# Correlating nuclear morphology and external force with combined atomic force microscopy and light sheet imaging separates roles of chromatin and lamin A/C in nuclear mechanics

**DOI:** 10.1101/2020.02.10.942581

**Authors:** Chad M. Hobson, Megan Kern, E. Timothy O’Brien, Andrew D. Stephens, Michael R. Falvo, Richard Superfine

## Abstract

Nuclei are constantly under external stress – be it during migration through tight constrictions or compressive pressure by the actin cap – and the mechanical properties of nuclei govern their subsequent deformations. Both altered mechanical properties of nuclei and abnormal nuclear morphologies are hallmarks of a variety of disease states. Little work, however, has been done to link specific changes in nuclear shape to external forces. Here, we utilize a combined atomic force microscope and light sheet microscope (AFM-LS) to show SKOV3 nuclei exhibit a two-regime force response that correlates with changes in nuclear volume and surface area, allowing us to develop an empirical model of nuclear deformation. Our technique further decouples the roles of chromatin and lamin A/C in compression, showing they separately resist changes in nuclear volume and surface area respectively; this insight was not previously accessible by Hertzian analysis. A two-material finite element model supports our conclusions. We also observed that chromatin decompaction leads to lower nuclear curvature under compression, which is important for maintaining nuclear compartmentalization and function. The demonstrated link between specific types of nuclear morphological change and applied force will allow researchers to better understand the stress on nuclei throughout various biological processes.

## Introduction

The nucleus – which encapsulates and protects the entire genome – functions not only as the site of gene replication and transcription, but also as a fundamental mechanical constituent of the cell. Altered nuclear mechanics and nuclear morphology have both been linked to various disease states ranging from Hutchinson-Gilford Progeria Syndrome (HGPS) (De Sandre-Giovannoli et al. 2003; Dahl et al. 2006; Butin-Israeli et al. 2012) and Emery-Dreifuss muscular dystrophy (Lammerding et al. 2004; Lammerding et al. 2005; Butin-Israeli et al. 2012) to breast cancer (Butin-Israeli et al. 2012; Tocco et al. 2018). Such diseased cells are constantly under stress either through external means such as cellular migration (Davidson et al. 2014; Harada et al. 2014) or intracellular forces like that of actin pre-stress (Lammerding and Wolf 2016), which has been shown to be sufficient to cause nuclear rupture (Hatch and Hetzer 2016; Lammerding and Wolf 2016). Little work, however, has studied either the dynamic relationship between external forces and nuclear morphology or the role of nuclear mechanical constituents in this relationship. In order to fully understand the complex connections linking nuclear mechanics and morphology with disease and cellular function, we must first understand the intermediate relationship of how nuclear mechanical constituents resists external forces to maintain morphology.

Nuclear mechanics are primarily dictated by the nuclear lamina and chromatin, as well as indirectly influenced by the cytoskeleton (Stephens, Banigan, and Marko 2019). The cytoskeleton protects the nucleus both through an actin cap (Khatau et al. 2009; Haase et al. 2016; Kim et al. 2018) and a peri-nuclear cage of the intermediate filament vimentin (Neelam et al. 2015; Patteson et al. 2019; Rosso, Liashkovich, and Shahin 2019). The nuclear lamina, primarily lamin A/C, has consistently been shown to be a major mechanical constituent of the nucleus through constricted migration, micropipette aspiration, atomic force microscopy, micromanipulation, and other techniques (Dahl et al. 2004; Lammerding et al. 2004; Dahl et al. 2005; Lammerding et al. 2006; Lee et al. 2007; Pajerowski et al. 2007; Schape et al. 2009; Swift et al. 2013; Hanson et al. 2015; Neelam et al. 2015; Stephens et al. 2017). Furthermore, understanding of chromatin’s role as a mechanical element of the nucleus continues to be refined. Through examining swollen *Xenopus* oocyte nuclei, it was first thought that chromatin had little role in the mechanical properties of nuclei (Dahl et al. 2004). Additional work, however, revealed that chromatin indeed does contribute to nuclear stiffness, and that (de)compaction of chromatin leads to nuclear (softening) stiffening (Dahl et al. 2005; Pajerowski et al. 2007; Mazumder et al. 2008; Krause, te Riet, and Wolf 2013; Erdel, Baum, and Rippe 2015; Schreiner et al. 2015; Shimamoto et al. 2017; Stephens et al. 2017; Neubert et al. 2018; Stephens et al. 2018). The specific roles of chromatin and lamin A/C in nuclear mechanics have begun to be disentangled, as micromanipulation experiments have shown that chromatin dominates small extensions while lamin A/C dominates large extensions (Stephens et al. 2017). Both the mechanical constituents of the nucleus – the nuclear lamina and chromatin – as well as the cytoskeleton are paramount for protection of the genome and subsequently cellular function.

Directly related to the mechanical properties of nuclei is nuclear morphology; this is in general characterized by nuclear volume and nuclear surface area – or the more experimentally accessible 2D surrogates of nuclear cross-sectional area and nuclear perimeter respectively – as well as local curvature. Nuclear morphology also relates to nuclear abnormalities and/or blebs displayed across the spectrum of human disease (Stephens, Banigan, and Marko 2019). However, here we are primarily concerned with morphology in regards to general nuclear shape. A variety of metrics have been used to quantify changes in nuclear morphology, such as area strain (percent change in projected cross-sectional area) (Zhang et al. 2019) and 3D irregularity (ratio of excess volume of a fitted convex hull to nuclear volume) (Tocco et al. 2018). Aside from the previously noted connections to disease, nuclear morphology has further been linked to levels of transcriptional activity as nuclei with reduced volume enter a more quiescent state (Damodaran et al. 2018). Increases in the volume of nuclei either through swelling (Finan, Leddy, and Guilak 2011) or directed migration on patterned substrates (Katiyar et al. 2019) has been shown to decondense or dilate chromatin levels. Stretching of the nuclear surface area is thought to be a mechanism of nuclear mechanotransduction (Enyedi and Niethammer 2017; Donnaloja et al. 2019). Nuclear morphology is also characterized in part by local curvature; regions of high local curvature have been linked to nuclear rupture (Xia et al. 2018) and nuclear blebs (Stephens et al. 2018; Cho et al. 2019). Nuclear morphology is directly related to both the mechanical integrity of the nucleus as well as nuclear and cellular function.

Previous work has used changes in nuclear morphology under force application as a metric for mechanical resistance (Neelam et al. 2015; Haase et al. 2016); that is, smaller changes in nuclear morphology imply a stiffer nucleus. Nuclear morphology has also been used in studying stored elastic energy (Tocco et al. 2018) and pressure gradients (Finan et al. 2009; Kim et al. 2015). Investigators have further developed an analytical model connecting nuclear morphology to external forces and mechanical properties for an idealized geometry (Balakrishnan et al. 2019). However, a majority of work regarding nuclear mechanics is either agnostic to nuclear shape or focuses on a highly specific model of a single technique. For example of the former, atomic force microscopy (AFM) studies of nuclei have traditionally used a Hertzian contact mechanics model, which models the nucleus as a linearly elastic, isotropic, homogeneous material under small indentation (Johnson 1985). Previous work, however, has shown the nucleus to be both nonlinear (Stephens et al. 2017) and anisotropic (Haase et al. 2016). While Hertzian analysis has brought to light many novel insights, it is limited by its ability to decouple contributions of specific structures. More intricate computational models have given direct insight into many mechanical techniques, including constricted migration (Cao et al. 2016), micropipette aspiration (Vaziri and Mofrad 2007), magnetic bead twisting (Karcher et al. 2003), plate compression (Caille et al. 2002), micromanipulation (Banigan, Stephens, and Marko 2017; Stephens et al. 2017), and atomic force microscopy (Lherbette et al. 2017); however, their specificity inhibits extrapolation of their conclusions. There exists a need for an intermediate understanding of nuclear deformation that informs both the relative contributions of the various nuclear mechanical constituents as well as their roles in protecting against specific deformations to nuclear morphology.

In this work, we address some of these open questions regarding the links between mechanics and morphology through use of our combined atomic force microscope and side-view light sheet microscope (AFM-LS) (Nelsen et al. 2019). Our approach allows us to visualize cells from the side (x-z cross section) with high spatiotemporal resolution during compression with an atomic force microscope. We use this technique to correlate changes in SKOV3 nuclear volume and nuclear surface area with applied force to develop an empirical model for nuclear deformation, which has applications for assays beyond our own technique and is applicable to non-standard nuclear shapes. This allows us to disentangle the contributions of chromatin and lamin A/C to strain in nuclear volume and nuclear surface area, respectively, an insight not possible with previous AFM models and techniques. We also measure the dynamics of nuclear curvature under compression, and show that chromatin decompaction reduces curvature at the site of indentation; this indirectly shows the nucleus behaves as a two-material system. To further interpret our findings, we develop a finite element analysis (FEA) model – allowing us to computationally study nuclear deformation for discretized, non-standard nuclear geometries – that recapitulates our empirical results and connects them to material properties. In summary, we provide the first decoupling of the role of chromatin and lamin A/C in nuclear compression as well as a new insight into the connection between external forces, nuclear mechanical constituents, and nuclear shape and curvature.

## Results

### Combined AFM and side-view light sheet microscopy show strain in nuclear volume and surface area begin at different indentations

Our combined atomic force microscope and side-view light sheet fluorescence microscope (Figure 1A) allows visualization of the dynamics of cellular deformations in the plane of applied force while simultaneously monitoring the force response of the cell (Nelsen et al. 2019). We have previously used this tool to show the existence of separate elastic moduli correlated with whole-cell and nuclear deformations (Beicker et al. 2018). We built on this previous work by studying the dynamics of nuclear morphology and the correlated force response under compression by AFM. We examined time series of side-view images of compressed, live SKOV3 cells stably expressing halotagged histone 2B (H2B, green) and snaptagged K-Ras-tail (magenta) labeled with Janelia Fluor (JF) 549 and 503 respectively (Figure 1B, Supplementary Movie 1). Masks of nuclei were generated (see Materials and Methods) and used to extract both nuclear cross-sectional area (*NCSA*, blue) and nuclear perimeter (*NP*, orange) as a function of indentation (Figure 1B). As in prior studies, we used *NCSA* and *NP* as surrogates for nuclear volume and nuclear surface area respectively as the qualitative deformation of the nucleus is the same in any side-view orientation (Finan, Leddy, and Guilak 2011). The AFM provided synchronized force data with approximately 20 pN resolution during the side-view image acquisition (Figure 1C, Supplementary Movie 1).

**Figure 1:**
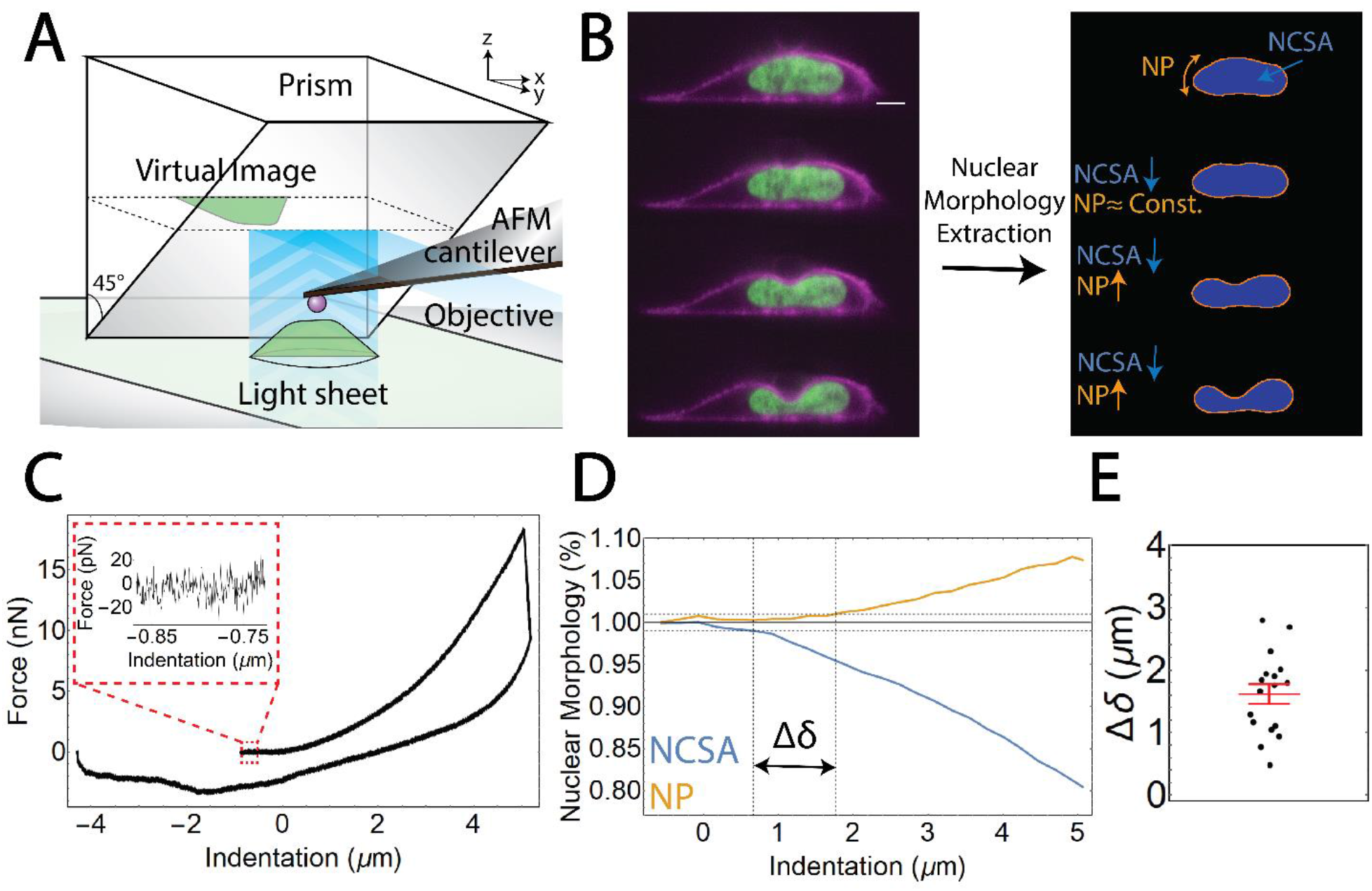
Combined atomic force microscopy and side-view light sheet microscopy (AFM-LS) extracts dynamics of nuclear morphology and applied force under whole-cell compression. (A) Cartoon schematic of our AFM-LS system. A full description is provided in our previous work. (B). A subset of fluorescence images collected by our AFM-LS during indentation of a live, SKOV3 cell (scale bar = 5 μm). The cell is stably expressing snaptagged KRas-tail (Magenta) and halotagged H2B (Green) labeled with Janelia Fluor 503 and 549 respectively. Custom workflow (see methods) allows for extraction of nuclear perimeter (*NP*) and nuclear cross-sectinoal area (*NCSA*). (C) Force versus indentation data for the previously displayed compression experiment. Inset provides a scale for the noise is the force data. (D) Nuclear morhpology as a percentage versus indentation for the previoysly displayed compression experiment. Orange and blue represent *NP* and *NCSA* respectively. Δ*δ* is defined to be the difference in indentation at which *NP* and *NCSA* reach 1% change. (E) Δ*δ* for n = 17 separate compression experiments. The red bar represents the standard error in the mean.

We first observed that *NCSA* and *NP* underwent strain at different levels of indentation (Figure 1D). We determined this difference in the onset of *NCSA* strain and *NP* strain by linearly interpolating the *NCSA* and *NP* indentation series and computing the difference in indentations at which *NCSA* and *NP* reached 1% strain, denoted by Δ*δ*. This was chosen because 1% strain is a point reached in all data sets used in analysis and is far enough above the noise of the nuclear morphology data to confidently indicate a change. The onset of strain in *NCSA* and *NP* differ by Δ*δ* = 1.6 ± 0.7 μm (mean ± standard deviation of indentation), which is clearly greater than zero. This indicates the presence of two distinct and separate regimes for strain onset of nuclear surface area and nuclear volume (Figure 1E).

### A two-regime force response allows for determination of scaling relationships between nuclear morphology and applied force

Previous research has shown both a two-regime force response upon stretching nuclei with flexible micropipettes (Stephens et al. 2017) as well as the necessity of a term accounting for the stretching of nuclear surface area to explain non-linear osmolarity of the nucleus (Finan et al. 2009). This work and our results showing distinct indentations thresholds for nuclear volume and surface area strain led us to hypothesize the existence of a two-regime force response resulting from separate forces associated with changes in nuclear volume and nuclear surface area.

To test this hypothesis, we first examined the scaling relationship between applied force from the AFM, *F*_AFM_, and *ΔNCSA* (*NCSA*_*Max*_ - *NCSA*); note that *ΔNCSA* is positive for a decrease in *NCSA*. We observed a clear, two-regime force response wherein applied force scales with *ΔNCSA* to different powers in each regime (Figure 2A). This phenomenon was seen in all but one cell examined (n=17 cells examined total). To determine the scaling relationships between external force and *ΔNCSA*, we fit two separate power law relationships between force and *ΔNCSA* – one before and one after the transition point. The transition point between fitting regimes was allowed to vary to minimize error in the power law fits in both regimes. The exact transition point was determined to be the point at which the two power law relationships intersect.

**Figure 2:**
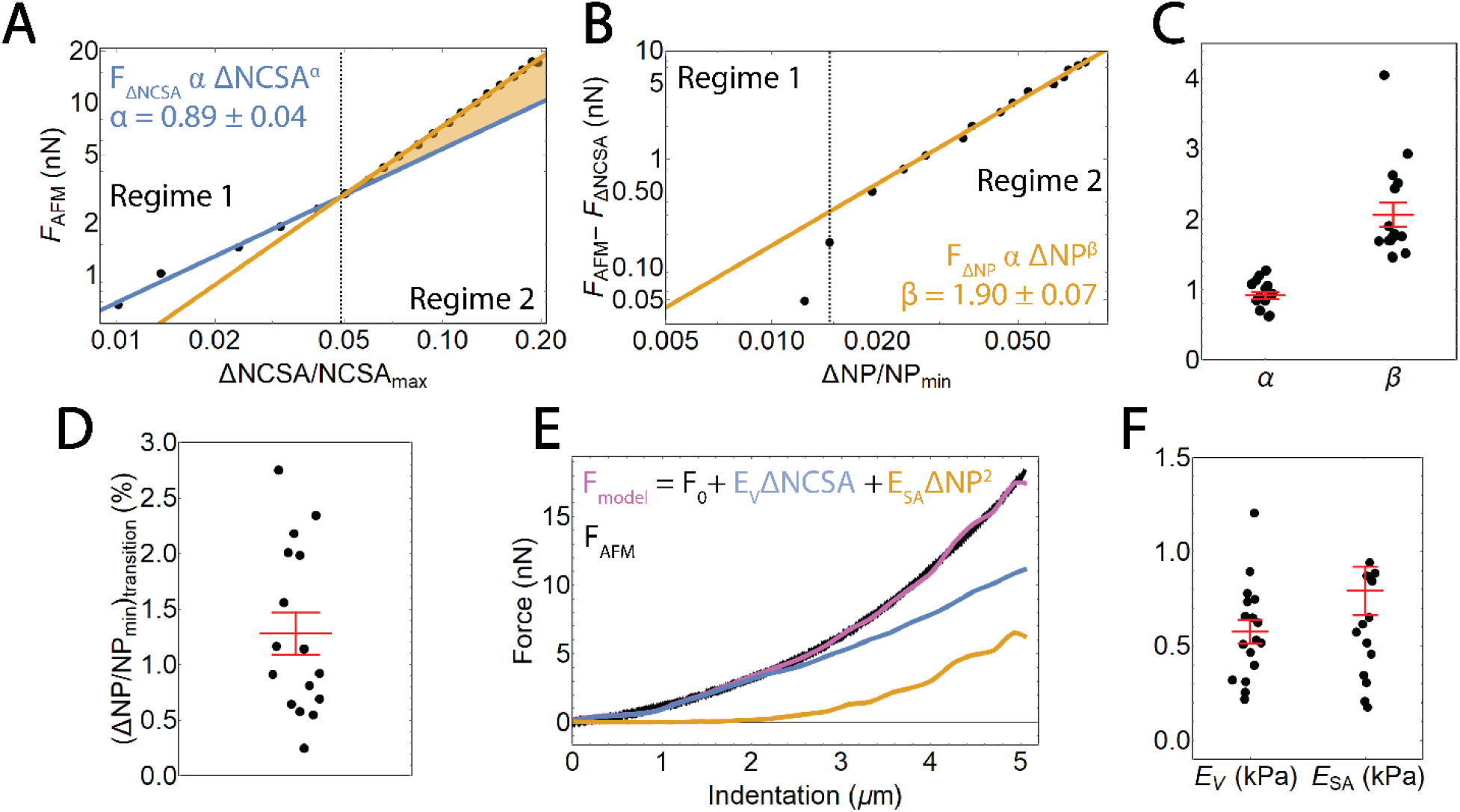
Correlating nuclear morphology and applied force informs an emperical model for strain stiffening response. (A) Force as recorded by the AFM versus change in *NCSA* plotted on a log-log scale. Two distinct power-law regimes are observed. (B) Force as recorded by the AFM minus the force response in regime 1 plotted against changed in NP on a log-log scale, showing a single power law relationship in regime 2. (C) α, the exponent for *F*_Δ*NCSA*_, as determinded for n and β, the exponent for *F*_Δ*NP*_, as determinded for n = 16 cells. Red bars represent mean and SEM. (D) The strain in *NP* at the transition point between regime 1 and regime 2 as determined for n = 16 cells. Red bars represent mean and SEM. (E) An emperical model for nuclear deformation as shown over force versus indentation. We display our full emperical model (magenta), the individual contributions required to deform the nuclear volume and surface area (blue, orange respectively), and the AFM data over the full indentaiton. (F) Resistance to nuclear volume change, *E*_V_, and resistance to nuclear surface area change, *E*_SA_, as determined for n = 17 cells. Red bars represent mean and SEM.

Knowing that at small indentations we only observed strain in *NCSA* indicates that regime 1 immediately provided us a scaling relationship between external force and *ΔNCSA*. That is, we defined a force associated with *ΔNCSA* given by *F*_Δ*NCSA*_ ∝ Δ*NCSA*^*α*^ (Blue line, Figure 2A). Under the assumption that the aforementioned relationship was unchanged during indentation, we subtracted *F*_Δ*NCSA*_ from *F*_AFM_ to isolate the additional force response resulting from strain in *NP* (Yellow shaded region, Figure 2A). We then plotted this additional force response against *ΔNP* (*NP – NP*_*Min*_) where we observed a constant power law relationship in regime 2 (Yellow line, Figure 2B). We then defined a separate force required to stretch the nuclear surface area given by *F*_Δ*NP*_ ∝ Δ*NP*^*β*^. Performing this analysis on n=16 cells allowed us to determine that *α* = 0.9 ± 0.2 and *β* = 2.1 ± 0.7 (mean ± standard deviation, Figure 2C).

Previous work, however, has modeled the nucleus as having a strain-dependent elastic modulus (Lherbette et al. 2017), which could provide an alternate explanation of the origin of the two-regime phenomenon we have observed. To differentiate the two explanations, we examined the transition point as a function of *ΔNP*. The transition point between the two regimes corresponded to 1.2% ± 0.8% (mean ± standard deviation) change in *NP* (Figure 2D), meaning the force response in regime 1 correlated only with strain in *NCSA* while the force response in regime 2 correlated with both strain in *NCSA* and *NP*. This correlation between the onset of regime 2 and the onset of strain in *ΔNP* provided support to our hypothesis that the two regimes are a result of separate forces required to deform the volume and surface area of the nucleus. With the combination of this result and our determination of the specific scaling relationships between applied force and both *ΔNCSA* and *ΔNP*, we then posed the following empirically-determined model to correlate nuclear deformation with applied force.

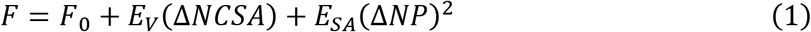

Here, *F*_0_ represents any force response accumulated prior to deformation of the nucleus, and *E*_*V*_ and *E*_*SA*_ are the effective mechanical resistance the cell provides to changes in nuclear volume and nuclear surface area respectively, as represented by *NCSA* and *NP*. These resistances are composed of contributions from not only the nucleus, but also from the cytosol, internal pressure gradients, the cytoskeleton, the actin cortex, and other cellular structures. However, the nucleus has been shown to be the stiffest sub-cellular structure and also encompasses a majority of the strain during compression, implying that *E*_*V*_ and *E*_*SA*_ are primarily dictated by the mechanical properties of the nucleus. Our results are consistent in that *F*_0_ is on the order of 100 pN, implying there is minimal force response prior to deformation of the nucleus. They also inherently include viscous contributions as there is no time scale built directly into our model and our AFM measurements are not fully quasistatic (Figure S1). Indenting at higher (lower) rates would then increase (decrease) our measured values of *E*_*V*_ and *E*_*SA*_. While not studied here, this decomposition provides the opportunity to study the relative viscous contributions associated with strain in *NCSA* and *NP*. A single value of *E*_*V*_ and *E*_*SA*_ are determined by fitting the Equation 1 to the entire indentation of each cell (Figure 2E and F).

We find that for SKOV3 cells, *E*_*SA*_ is approximately 1.36 times greater than *E*_*V*_, implying these nuclei are more susceptible to strain in volume than in surface area. This can be compared to the Hertz model (Johnson 1985) and the height-corrected Hertz model (Dimitriadis et al. 2002), both of which fail to model the force response over the entirety of the indentation (Figure S2). It is also important to note that Hertzian analysis requires small indentations, specific probe and target geometries, and assumptions regarding homogeneity and linearity. Our approach, however, makes no prior assumption regarding such geometries and is simply empirical. Furthermore, our approach allows us to decouple resistances to specific types of nuclear strain as opposed to providing a single metric of stiffness for the entirety of the nucleus. This complements and improves on earlier analytic modeling efforts (Balakrishnan et al. 2019) in that we have empirically determined a relationship between force and morphology that accounts for contributions of both the bulk compressibility and surface tension without assuming a predefined geometry.

### Empirical model of nuclear deformation is independent of initial nuclear size

Our AFM-LS technique does not systematically examine a specifically oriented vertical slice as the distribution of polarity amongst the cells is seemingly random. One potential failure of the proposed model and technique would be a dependence of *E*_*V*_ and *E*_*SA*_ on the initial morphology of the nucleus or the orientation in which we image the nucleus from the side. To examine this, we determined both *E*_*NCSA*_ and *E*_*NP*_ for n = 17 cells and plotted *E*_*V*_ and *E*_*SA*_ against initial values of *NCSA* and *NP* (Figure S3). We performed a Pearson’s correlation test between *E*_*V*_, *E*_*SA*_ and *NCSA*, *NP*. No significant correlation was observed between either *E*_*V*_ or *E*_*SA*_ and *NCSA* or *NP*, implying that the resistances to nuclear morphology changes determined by our model are not systematically dependent on either the scale of the nucleus or the specific side-view orientation in which we visualized the nucleus. Our approach is then robust to any initial cell orientation or initial nuclear size.

### Chromatin and lamin A/C separately resist nuclear volume and surface deformations respectively

Chromatin and lamin A/C have been shown to be the primary mechanical constituents of nuclei; recent work has shown that during micromanipulation extension of isolated nuclei chromatin dominates small-scale extensions while lamin A/C governs large-scale extensions (Stephens et al. 2017). It remains untested if similar phenomena hold true for compression-based deformations of nuclei in intact cells. We hypothesized that in AFM indentations chromatin in part dictates the resistance to nuclear volume change while lamin A/C separately resists changes in nuclear surface area. Such a measurement was not previously attainable without AFM-LS.

To test this hypothesis, we first treated our SKOV3 cells with a 200 nM concentration of Trichostatin A (TSA) for 24 hours prior to performing AFM compression with side-view imaging experiments. TSA decompacts chromatin by increasing euchromatin marker histone tail acetylation (Figure S4) (Toth et al. 2004). We extracted nuclear morphology dynamics under compression and fit Equation 1 to the corresponding force data to extract *E*_*V*_ and *E*_*SA*_. We observed a significant 40% decrease in *E*_*V*_ upon TSA treatment (p<0.05 from a T Test), but no significant difference in *E*_*SA*_ (Figure 3A and B).

**Figure 3:**
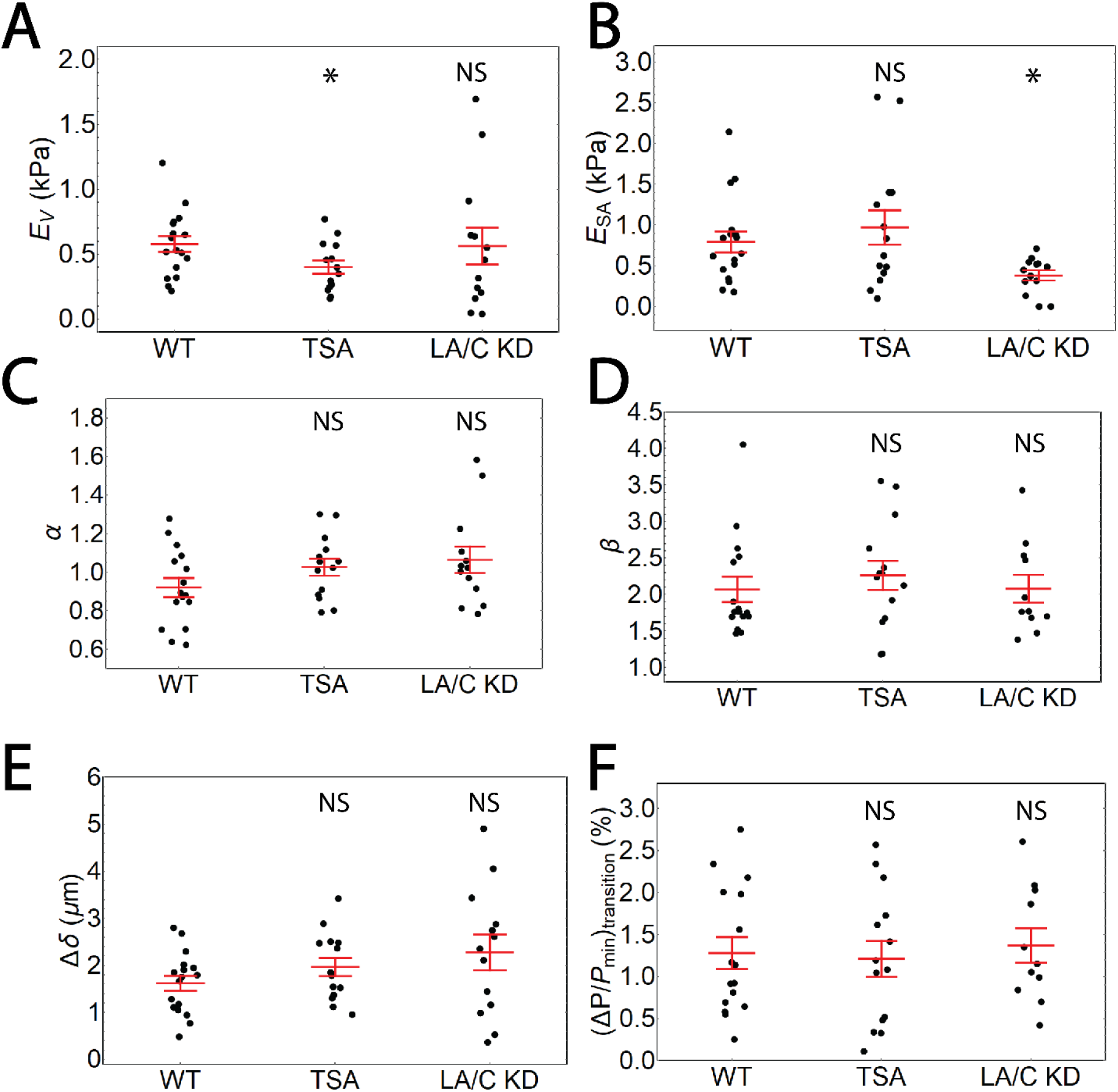
Chromatin decompaction and lamin A/C depletion weaken resistance to volume and surface area changes respectively, while behaving similar to the empirical model. (A) Resistance to nuclear volume change, *E*_V_, is decreased by TSA but unchanged by LA/C KD. n = 17, 14, 13 for WT, TSA, LA/C KD. (B) Resistance to nuclear surface area change, *E*_V_, is unchanged by TSA but decreased by LA/C KD. n = 17, 14, 13 for WT, TSA, LA/C KD. (C) α, the exponent for *F*_Δ*NCSA*_, is unchanged by TSA and LA/C KD. n = 16, 14, 13 for WT, TSA, LA/C KD. (D) β, the exponent for *F*_Δ*NP*_, is unchanged by TSA and LA/C KD. n = 16, 14, 11 for WT, TSA, LA/C KD. (E) The difference in indentation at the onset of change in NP and NCSA is unchanged by TSA and LA/C KD. n = 17, 14, 13 for WT, TSA, LA/C KD. (F) The strain in NP at the transition point between regime 1 and regime 2 is unchanged by TSA and LA/C KD. n = 16, 14, 11 for WT, TSA, LA/C KD. Red bars represent mean and SEM. NS – not significant. * - p < 0.05.

Furthermore, we transfected our SKOV3 cell line with siRNA to halt production of new lamin A/C (LA/C KD, Figure S5). We then performed AFM-LS experiments 4-6 days post transfection and extracted *E*_*V*_ and *E*_*SA*_ as previously described. We observed a significant 50% decrease in *E*_*SA*_ (p<0.05 for a T Test), yet no significant change in *E*_*V*_ (Figure 3A and B). This means both that chromatin resists strain in nuclear volume while lamin A/C separately resist strain in surface area. Furthermore, this indicates that chromatin does not resist nuclear surface area stretching nor does lamin A/C resist deformation in nuclear volume. Because strain in nuclear volume and nuclear surface area occur at different indentation scales (Figure 1), we have also shown that chromatin and lamin A/C provide mechanical resistance at short and long indentations respectively.

### Empirical model of nuclear deformation is independent of modifications to chromatin and Lamin A/C

An alternative explanation for the decreases in *E*_*V*_ and *E*_*SA*_ seen upon TSA treatment and LA/C KD respectively is that our proposed empirical model (Equation 1) is no longer valid after these treatments. More specifically, we could be observing changes in *E*_*V*_ and *E*_*SA*_ that are actually representative of changes in the scaling relationships between force and nuclear morphology themselves; that is, *α* and *β* could be dependent on chromatin compaction and lamin A/C expression. To address is explanation, we performed the analysis previously described to extract *α* and *β* from both the TSA-treated and LA/C KD samples. We found no significant change in either *α* or *β* (Figure 3C and D), meaning the previously determined scaling relationships between nuclear morphology and applied force are unchanged. Because these scaling relationships remain constant, our observed changes in *E*_*V*_ and *E*_*SA*_ are indicative of changes in the nucleus’ ability to resist strain in nuclear volume and nuclear surface area.

Furthermore, we observed no significant difference in Δ*δ* (Figure 3E) or the strain in *NP* at the transition point (Figure 3F). This implies that the existence of the strain-stiffening effect is more closely related to nuclear geometry and the manner of deformation than the relative stiffnesses associated with the volume and surface area of the nucleus. This result is supported by previous findings in micromanipulation studies (Banigan, Stephens, and Marko 2017). The lack of changes in scaling (*α* and *β*) and transition (indentation and strain) solidifies our conclusions regarding the role of chromatin and lamin A/C in separately resisting strain in nuclear volume and surface area respectively.

### Chromatin decompaction and Lamin A/C KD separately regulate nuclear curvature under compression

Recent studies have connected nuclear curvature to locations of nuclear rupture and subsequent DNA damage. Through AFM compression with both 4.5 μm diameter beaded cantilevers and sharp tip cantilevers (diameter < 0.1 μm), a correlation between indentation with high-curvature probes and nuclear rupture was reported (Xia et al. 2018). Similarly, nuclear blebs induced by increases of euchromatin were shown to systematically form at the pole of the major axis, which is the site of highest curvature (Stephens et al. 2018). We then sought to study the roles of chromatin and lamin A/C in nuclear curvature dynamics during AFM compression.

Nuclear curvature, defined as the inverse of the radius of a best-fit circle, was extracted for each point in the discretized perimeter of the nucleus for every image collected during the indentation. Prior to compression, the nucleus shows two peaks corresponding to the ends of the oval-shaped nucleus (Figure 4A). Once compressed, the nucleus shows a new, clearly defined peak at approximately 10 μm; this peak corresponds to the new curvature formed as a result of indentation with the AFM (Figure 4B, Supplementary Movie 2). By fitting a Gaussian curve to this peak for each frame in the indentation (Figure 4C), we can study how curvature changes as a function of indentation; specifically, we extract maximum curvature at the site of indentation. We observed that in the regime over which nuclear curvature changes, there is a linear relationship between maximum nuclear curvature and indentation (Figure 4D). Eventually the maximum nuclear curvature plateaus as it cannot exceed that of the AFM probe.

**Figure 4:**
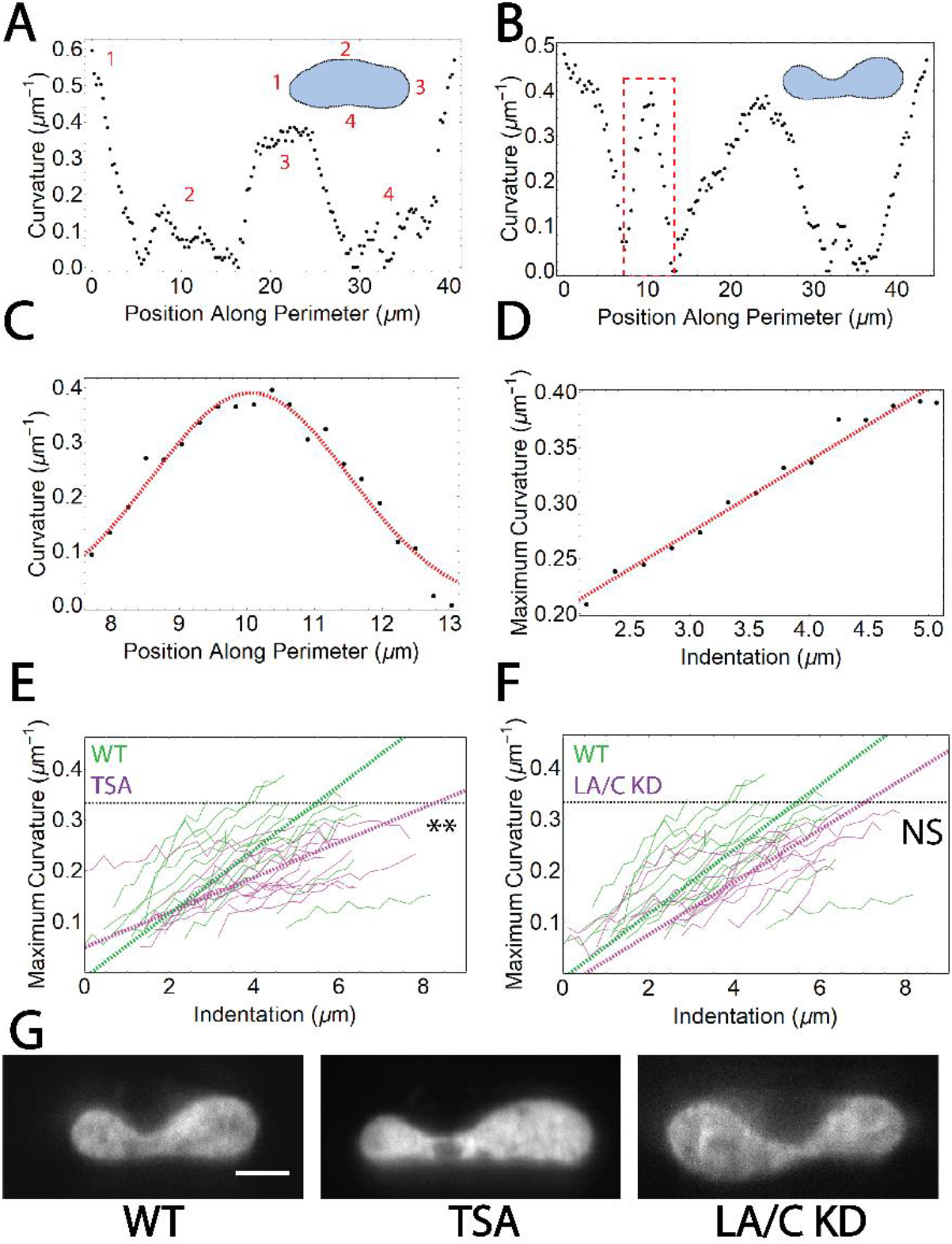
Dynamic nuclear curvature analysis during AFM indentation. (A) Nuclear curvature versus position along perimeter for an undeformed nucleus. The inset displays the mask of the nucleus and the discritization of the perimeter. (B) Nuclear curvature versus position along perimeter for a deformed nucleus. The inset displays the mask of the nucleus and the discretization of the perimeter. Note the additional peak centered around 10 μm representing the site of indentation. (C) A Gaussian fit to the peak at the site of indentation extracts maximum curvature under the AFM bead (red dashed box in (B)). (D) Maximum curvature plotted during the entire indentation. A linear fit is performed for the region of changing curvature. (E) Maximum curvature as a function of indentation plotted for n = WT cells and n = TSA treated cells. ** represnts p < 0.01 for a T Test comparing the mean slope of maximum curvature versus indentation. (F) Maximum curvature as a function of indentation plotted for n = WT cells and n = LA/C KD cells. Black dashed line represents curvature of the AFM bead. NS represnts no significance for a T Test comparing the mean slope of maximum curvature versus indentation. (G) Representative images of SKOV3 nuclei (H2B) under maximum compression with various treatments. Scale bar = 5 μm.

Maximum nuclear curvature was plotted against indentation and compared between perturbation and control WT nuclei (Figure 4E-G). We discovered a significant decrease (p<0.01 for a T Test) between the mean slope of maximum curvature versus indentation for TSA treated cells. This implies that larger indentations are necessary to induce that same of amount of nuclear curvature in TSA treated cells as compared to WT cells. Contrary to chromatin decompacted cells, we observed no significant change in the mean slope of maximum curvature versus indentation for LA/C KD cells as compared to WT cells; this could be due to either a minimal contribution of lamin A/C to nuclear curvature or an inability to detect the changes in nuclear curvature due to the geometry of our assay or the degree of knockdown of lamin A/C. While not studied here, our ability to monitor dynamics of nuclear curvature under compression will facilitate further studies of both varied deformation geometries and increased of chromatin compaction and lamin A/C levels.

### Two-material finite element model correlates resistances to morphology changes with material properties

In order to validate our conclusions and connect our results to mechanical properties, we developed a simple computational model of AFM indentation experiments. As the nucleus is the stiffest sub-cellular structure and the focus of our analysis, we chose to model only deformation of the nucleus. We constructed an axisymmetric finite element analysis (FEA) model featuring a stiff, spherical, polystyrene indenter and an ellipsoidal nucleus (Figure 5A, Movie S3). Previous research has examined FEA models of AFM (Chen and Lu 2012; Liu, Mollaeian, and Ren 2019), yet to our knowledge none have examined the relationship of nuclear morphology and force. The nucleus in our model features two separate materials: an infinitely thin, elastic membrane with a stretch modulus (*γ*) wrapped around an elastic solid with an elastic modulus (*E*). The model assumes quasistatic behavior.

**Figure 5:**
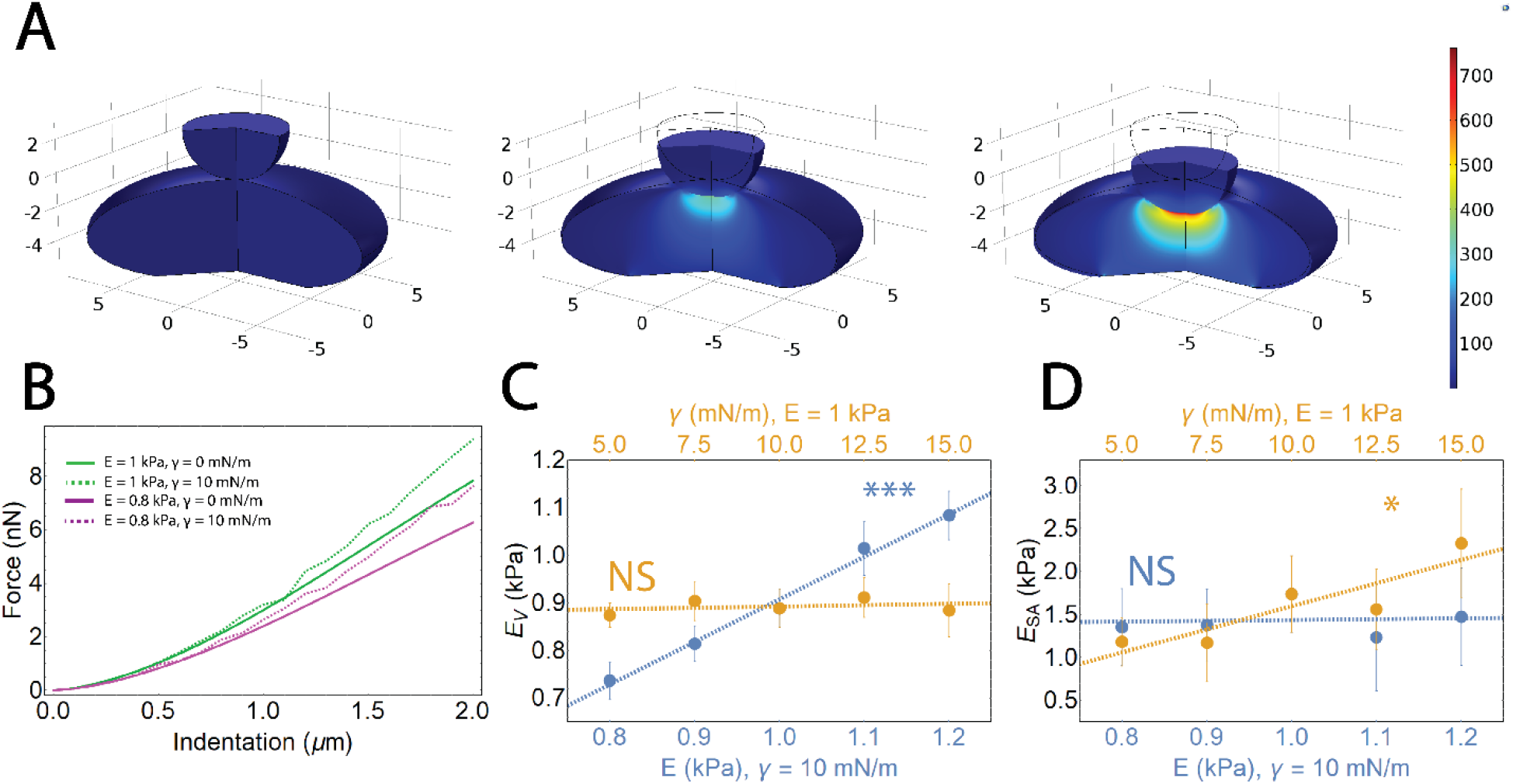
Finite Element Analysis (FEA) model of AFM indentation. (A) Selected frames from a finite element analysis simulation of a nucleus undercompression. The nucleus has an elastic modulus, *E*, and a separate stretch modulus, *γ*. (B). Force versus indentation shown for varied *E* and *γ*. (C). Resistance to nuclear volume change, *E*_*V*_, plotted against variations in both *E* and *γ*. A significant correlation (p<0.001) between *E*_*V*_ and *E*, but no significant correlation is seen between *E*_*V*_ and *γ*. (D)). Resistance to nuclear surface area change, *E*_*SA*_, plotted against variations in both E and *γ*. A significant correlation (p<0.05) between *E*_*SA*_ and *γ*, but no significant correlation is seen between *E*_*V*_ and *E*.

We simulated AFM indentation with varied values of *E* and *γ* and examined the qualitative changes in the force-indentation curves (Figure 5B). We found that varying the elastic modulus leads to softening over the entirety of the indentation. However, variations in the stretch modulus of the elastic layer led to altered behavior only at larger indentations (> 0.5 μm). This corresponds directly with our observation from our AFM-LS experiments. We showed that decondensation of chromatin and knockdown of lamin A/C lead to decreases in *E*_*V*_ and *E*_*SA*_ respectively, and *E*_*V*_ provides resistance over the entire indentation while *E*_*SA*_ provides the additional resistance at larger indentations.

Observing the same qualitative behavior in the force-indentation curves provided motivation to correlate the material properties of the FEA model (*E*, *γ*) with *E*_*V*_ and *E*_*SA*_. To do so, we performed identical analysis for extracting *E*_*V*_ and *E*_*SA*_ as previously described on our FEA model data. By varying either *E* or *γ* while keeping the other constant, we were able to study these correlations. We examined correlations between *E*_*V*_ (Figure 5C) and *E*_*SA*_ (Figure 5D) with *E* (blue) and *γ* (orange). We observed a significant, linear correlation between *E*_*V*_ and *E* as well as *E*_*SA*_ and *γ*; no significant correlation was observed between either *E*_*V*_ and *γ* or *E*_*SA*_ and *E*. This shows that our resistances to nuclear volume change and nuclear surface area change (*E*_*V*_ and *E*_*SA*_) are indicative of material properties of the nucleus. Specifically, *E*_*V*_ is a representative measure of the elastic modulus of the nucleus and *E*_*SA*_ is a representative measure of the nuclear stretch modulus. We then find that our measured mean *E*_*V*_ of 0.58 kPa corresponds to *E* = 0.63 kPa and our measured mean *E*_*SA*_ of 0.79 kPa corresponds to *γ* = 2.5 mN/m, both of which are consistent with the current literature for Young’s modulus measurements of the nucleus and stretch modulus of the nuclear membrane respectively (Dahl et al. 2004; Dahl et al. 2005; Mazumder et al. 2008; Schape et al. 2009; Liu et al. 2014; Neubert et al. 2018; Wang et al. 2018; Rosso, Liashkovich, and Shahin 2019). Furthermore, we varied the size of our model nucleus and found that *E*_*V*_ and *E*_*SA*_ are independent of initial nuclear size (Figure S6), similar to our results from experiments (Figure S3).

## Discussion

### Nuclear morphology provides an estimate of external forces

Our combined AFM-LS approach to studying nuclear mechanics first showed the existence of two regimes of deformation: one at low levels of indentation (regime 1) and one at high levels of indentation (regime 2). In regime 1, we observed only changes in nuclear volume; in regime 2, we observed changes in both nuclear volume and surface area. This allowed us to extract scaling relationships between applied external forces and changes in nuclear morphology, leading to an empirical model of nuclear deformation characterized by two fitting parameters: *E*_*V*_ and *E*_*SA*_ (Figure 2). These fitting parameters provide a metric of resistance to nuclear volume change and nuclear surface area change, respectively, but further are directly proportional to the elastic modulus of the nucleus and the nuclear stretch modulus (Figure 5).

Often missing from current studies, are measurements of the external stress a nucleus experiences throughout a given experiment. Such measurements could provide additional context for interpreting why certain phenomena may be occurring. For example, constricted migration assays how allowed investigators to study how deficiencies in the nuclear lamina lead to increased migration rates (Shin et al. 2013; Davidson et al. 2014; Harada et al. 2014), higher rates of nuclear rupture (Denais et al. 2016), and increases in plastic damage (Harada et al. 2014; Davidson et al. 2015; Cao et al. 2016), all of which are relevant for understanding disease states and cellular function. Our work provides a means of estimating the external force applied to a nucleus simply through measuring nuclear morphology in the plane of applied force. Clear limitations exist for applying this model, such as complex 3D nuclear strains; however, some common assays for studying nuclear dynamics could benefit for our model.

### Chromatin and lamin A/C govern different forms of deformation

With our approach we were able to separate deformations of nuclear volume and nuclear surface area and tease apart how the mechanical constituents of the nucleus are responsible for each class of deformation. Specifically, we showed through disruption of histone-histone interactions via TSA treatment that chromatin resists strain in nuclear volume, but not in the nuclear surface area. We also showed through a knockdown of lamin A/C that the nuclear lamina resists strain in the nuclear surface area, but not in the nuclear volume. This further implies that chromatin dictates the mechanics of small indentations, whereas lamin A/C is relevant for large indentations. Additionally, through our FEA modeling we find that alterations in chromatin compaction and lamin A/C expression directly alter the nuclear elastic and stretch moduli respectively. These findings are in agreement with work showing that stretching of isolated nuclei via micromanipulation with micropipettes yields a two-regime force response where chromatin regulates the low-deformation regime and lamin A/C regulates the high-deformation regime (Stephens et al. 2017). These results help to provide a guide for AFM experiments in that small indentation measurements will likely probe only the mechanical properties of the chromatin, and large indentation measurements will need additional compensation beyond standard contact mechanics models to account for stretching of the nuclear lamina.

Previous research has also shown that in micropipette aspiration studies, the mechanics of swollen nuclei are dominated by the nuclear lamina whereas the mechanics of shrunken nuclei are governed in part by chromatin (Dahl et al. 2005). This result that shifting the primary load-bearing structure from chromatin to the nuclear lamina via swelling is consistent with our results as swelling would pre-stretch the nuclear surface area, thus eliminating regime 1 entirely and leaving only the lamina-dominated regime 2. Studies of osmolarity have also shown similar phenomena that can be explained by our model. Cell volume and inverse osmolarity follow a linear relationship, which can be modeled by the Boyle Van’t Hoff relation (Nobel 1969). Nuclei, however, have been shown to deviate from this linear relationship at large swelling volumes. To match this behavior, an additional term modeling nuclear membrane tension which is proportional to the nuclear surface area must be added (Finan et al. 2009), consistent with our findings.

AFM studies of mechanical properties have historically been used to study small-deformation mechanical properties because of the limitations of the analytical models applied to resulting data. Using cantilevers with large (10 μm) diameter beads positioned above the nucleus, force-indentation curves were collected on HT1080 fibrosarcoma cells for relatively low indentations (~1 μm). The Hertz model was fit to the force-indentation curves to extract an elastic modulus. They showed that TSA treatment resulted in nuclear softening (Krause, te Riet, and Wolf 2013). This is consistent with our results showing that TSA treatment reduces *E*_*V*_, which is the primary resistance at low indentations and proportional to the elastic modulus of the nucleus. A separate AFM study on isolated *Xenopus* oocytes revealed not only that increases in lamin A resulted in nuclear stiffening, but that the force-indentation curves become more linear as a function of lamin A expression levels (Schape et al. 2009); a linear force-indentation curve is representative of a pressurized shell (Vella et al. 2012). They were able to extract a nuclear membrane tension by fitting the end of the force indentation curve to a linear relationship, and it is crucial to note that they indented nuclei up to approximately 10 μm. By fitting the end of the force-indentation curves for large indentations, they effectively extracted the mechanical properties of regime 2. Our results are then consistent in that we also showed a dependency of *E*_*SA*_ on lamin A/C expression; this implies that lower levels of lamin A/C reduces the nuclear stretch modulus at large nuclear deformations.

Our work provides a synthesis of these previous studies, helping to piece together separate results into a single, cohesive explanation for the force response of the nucleus. Previous AFM studies using Hertzian analysis could only observe when nuclei became softer or stiffer under a given treatment. That is, chromatin decondensation and lamin A/C knockdowns show an apparently identical softening response (decrease in elastic modulus) under Hertzian analysis (Krause, te Riet, and Wolf 2013; Rauschert et al. 2017), despite having distinctly different roles in nuclear compression as shown here. With knowledge of both the length scales of deformation for which chromatin and lamin A/C are relevant as well as the specific morphological deformations against which they protect, we can progress forward to understanding their relative contributions in disease and function.

### The role of nuclear curvature

We observed that chromatin decondensation through TSA treatment resulted in less nuclear curvature during indentation as compared to WT, whereas LA/C KD nuclei showed no change in curvature under compression in our assay. We hypothesize that this is due to a change in the relative resistances to nuclear volume and nuclear surface area strains that we have previously shown. By decreasing the bulk resistance, the energy necessary to deform the nuclear volume relative to the nuclear surface area decreases. Minimization of energy cost then implies that the nucleus will undergo larger volume changes and smaller surface area changes. Decreased nuclear curvature is one means by which the nucleus could accommodate larger volume changes with less stretching of the nuclear surface area. Our hypothesis for the decreased rate of curvature change in TSA treated cells would lead us hypothesize that LA/C KD cells would show a higher rate of curvature change, or that overexpression of lamin A/C would mimic the behavior of TSA treated nuclei. Our observation of no significant change could either be due to a lack of correlation between lamin A/C and nuclear curvature, or more likely due to the geometry of the AFM probe and minimal knockdown of lamin A/C levels limit our ability to observe the affect LA/C KD would have on nuclear curvature. Regardless, the dependence of nuclear curvature on chromatin compaction levels clearly shows that the nucleus behaves as a two-material system. For a simple elastic solid, the curvature at the site of indentation is independent of the Young’s modulus, a second material is necessary to observe altered deformation patterns by changing the material properties. This further confirms that both chromatin and lamin A/C contribute to nuclear stiffness in compression.

The relevance of nuclear curvature has primarily been linked to rupture of the nuclear envelope as well as development of nuclear blebs. Previous work has examined the correlation between nuclear curvature and nuclear envelope rupture through AFM (Xia et al. 2018). U2OS cells were compressed with a constant force using both sharp (diameter < 0.1 μm) tips and 4.5 μm beads. Nuclear rupture was shown to be significantly more frequent when using the sharp tip as detected by mislocalization of YFP-NLS into the cytoplasm. Similarly, a constricted migration assay has revealed that it is rather the nuclear curvature, as opposed to tension, that is relevant for nuclear envelope rupture (Xia et al. 2019). Nuclear blebs have also been shown to systematically form at sites of high curvature (Stephens et al. 2018; Cho et al. 2019) and are prone to rupture (Stephens et al. 2019). Specifically, chromatin decompaction alone was sufficient to induce an increase in nuclear blebs (Stephens et al. 2018). Nuclear curvature is then highly relevant for understanding the mechanical integrity of nuclei, as loss of nuclear compartmentalization due to rupture causes nuclear dysfunction which may contribute to human disease (Davidson and Lammerding 2014; Stephens, Banigan, and Marko 2019). Our work suggests that lamin A/C may not be important for nuclear curvature. However, a valid alternate hypothesis is that the geometry of our bead limits our ability to detect the effect of the lamin A/C KD on nuclear curvature. We have clearly shown, however, that the state of compaction of chromatin has a direct link to the nuclear curvature, and we hypothesize that this is due to the altered, relative contributions of the nuclear elastic modulus and nuclear stretch modulus.

Our results regarding the dynamics of nuclear curvature under indentation also provide insight into how nuclear mechanotransduction may be altered through chromatin decompaction. We show that the nucleus develops less curvature during indentation for TSA treated cells, simultaneously implying that there is less stretching of the nuclear envelope. As previously noted, stretching of the nuclear lamina is thought to be a fundamental mechanism of nuclear mechnotransduction (Enyedi and Niethammer 2017). The state of compaction of chromatin may then indirectly alter transcription or the function of stretch activated channels (Donnaloja et al. 2019) if the nucleus is undergoing external stress, as we have shown the distribution of strain to be dependent on chromatin compaction.

### Nuclear morphology and function

Nuclear volume has been shown to be directly correlated with transcriptional activity. In one study, NIH3T3 mouse fibroblasts cultured on fibronectin patterned coverslips were uniformly compressed with an additional weighted coverslip. This resulted both in an increase in chromatin condensation as well as a decrease in nuclear volume, both of which correlated with the subsequent reduction in transcriptional activity (Damodaran et al. 2018). Separately, a migration assay has been used to show that transcription activity is altered as a result of introducing a constriction to the migration pathway (Jacobson et al. 2018), which can decrease nuclear volume. Through showing chromatin is partially responsible for resisting nuclear volume strain and coupling this result with previous studies regarding nuclear volume and function, we conjecture that the mechanical properties of chromatin aid in regulating its own condensation and transcriptional activity. We further see that the nucleus is susceptible to volume changes at low levels of indentation, meaning these downstream effects of volume change can occur as a result of intracellular forces.

Stretching of the nuclear surface, however, has different implications for nuclear function and mechanotransduction (Enyedi and Niethammer 2017). The nucleus is a mechanosensor that can covert mechanical signals at the cell surface into chemical responses (Kirby and Lammerding 2018). This physical connection from integrins to the nucleus through the cytoskeleton, the LINC complex, and the nucleus was first shown by pulling fibronectin-coated micropipettes attached to the cell surface (Maniotis, Chen, and Ingber 1997). It was later shown that by twisting fibronectin-coated magnetic beads attached to the cell surface, one could induce stretching of chromatin and subsequent upregulation of transcription activity (Tajik et al. 2016). The distribution of stresses along the nuclear lamina is believed to be the primary mechanism responsible for such responses (Enyedi and Niethammer 2017), which has led to the hypothesis of stretch-activation along the nuclear envelope (Donnaloja et al. 2019). By connecting expression levels of lamin A/C to resistances to change in nuclear surface area and consequently the nuclear stretch modulus, we have shown the relevance of lamin A/C to mechanoresponses governed by stretches in the nuclear envelope. Interestingly, we observe such nuclear surface area stretches only at large deformations. This implies that the nuclear lamina is relevant primarily for processes such as cellular migration or joint compression, and that the mechanoresponses associated with nuclear envelope stretches are not likely to happen outside of such processes that cause macroscopic, whole-cell deformations.

## Materials and Methods

### Plasmid construction

LZ10 PBREBAC-H2BHalo was a gift from James Zhe Liu (Addgene plasmid #91564; http://n2t.net/addgene:91564; RRID: Addgene_91564) (Li et al. 2016). pR-pre-EGFP was a gift from Sergio Grinstein (Addgene plasmid # 17274; http://n2t.net/addgene:17274; RRID: Addgene_17274) (Yeung et al. 2006). Piggybac plasmid PB513Bm2 was made by removing copGFP from PB513B-1 (System Biosciences) by PCR-based mutagenesis. PBREBAC_H2BHalo and PB513Bm2 encode the G418 and puromycin resistant gene, respectively. The PB513Bm2_SNAP-KRas-tail vector was generated using PB513Bm2 as the backbone and SNAP_KRas-tail as the insert. Specifically, EGFP in pR-pre-EGFP was replaced with SNAP tag, then both pR-pre_SNAP_KRas-tail and PB513Bm2 were digested using NheI-HF and BamH1-HF restriction enzymes (New England Biolab, NEB). The products were purified using the QiaQUICK Gel Extraction Kit protocol (Qiagen) and then ligated together using T4 DNA ligase (NEB) according to the manufacturer’s instructions. Prior to use, the plasmid sequences were confirmed by sequencing using the CMV-Forward primer at Genewiz (NJ).

### Generation of SKOV3 cell line and cell culture

SKOV3 cells were obtained from ATCC (HTB-77) and maintained in DMEM (Corning 15013CV) supplemented with 10% FBS (Sigma-Aldrich) and 1% GlutaMAX (Gibco). SKOV3 cells co-expressing H2B-Halo and SNAP_KRas-tail were generated through two consecutive transfections. We first produced a stable SKOV3 cell line expressing H2B-Halo to label the nucleus and then used those cells to produce a stable cell line expressing SNAP_KRas-tail to label the plasma membrane. The transfection was performed using Fugene HD transfection reagent (Promega) according to manufacturer’s instructions. Briefly, SKOV3 cells were seeded onto a 6-well plate at 5×10^4 cells per well 24 hours prior to transfection. For each well, we used 6 ul of Fugene HD transfection reagent, 3 μg piggyBac transgene plasmid (LZ10 PBREBAC_H2BHalo or PB513Bm2_SNAP_KRastail) and 0.6 μg of piggyBac transposase plasmid (ratio at 5:1). After 24 hours transfection, the medium was replaced with the fresh culture medium and the cells were recovered for 24 hours. Stable transfectants were selected by gradually increasing antibiotics concentrations; geneticin (G418) at a concentration to 1 mg/ml for PB-Halo-H2B and Puromycin at a concentration to 2 μg/ml for PB513Bm2-SNAP-KRas-tail. SKOV3 cells were grown in DMEM F12 without phenol red (Gibco), 5% FBS (Sigma-Aldrich) and 1X antibiotic antimycotic (Gibco). On the day before experiments they were trypsinized and plated at low density on fibronectin-coated polyacrylamide gels. 10 μL of Janelia Fluor 549 and 503 was added 2 hours prior to experiments, and washed out immediately before cell were examined. Janelia Fluor 503 was not used in Lamin A/C KD cells as to not conflate the GFP reporter signal.

### Production of polyacrylamide gels

Polyacrylamide gels were used as the cell substrate in order to eliminate reflections during side-view imaging. They were therefore produced to be at high stiffness (55 kPa), and relatively thin (10-30 um thick). They were produced and coated with fibronectin by the methods described in our previous work (Nelsen et al. 2019). Briefly, 10 μL of activated gel solution was deposited on APTES-treated 40 mm round coverslips, and a 22×22 mm square coverslip quickly placed on top. The top coverslip had been treated with HMDS via vapor deposition to facilitate easy removal after polymerization, and the gel included 1% polyacrylacrylic acid to provide carboxylic acid groups within the gel and promote adhesion to the APTES coated glass substrate. After gelation and coverslip removal under deionized water, the gel was allowed to dry in a Biosafety hood, sufficiently to allow placement of a 10 mm diameter glass cloning cylinder (316610, Corning) lightly coated with vacuum grease (1597418, Dow Corning). As soon as each cloning cylinder was placed, a solution of 10 mg/mL EDAC and 1 mg/mL NHS in PBS was placed into the cloning rings. The assembly was placed into sterile plastic petri dishes. The dishes were then placed in a 37°C chamber at 100% humidity for 15 minutes. The EDAC buffer was then replaced twice with PBS at room temperature, and then with 10 μg/mL fibronectin for 30 minutes at 37°C. The fibronectin solution was replaced with PBS twice, and then with DMEM F12 growth media. Samples were finally placed in the cell culture incubator to equilibrate at least 30 min before cells are added.

### Combined atomic force microscopy and side-view light sheet microscopy

The intricate details our AFM-LS system, both regarding the optical design and integration of the atomic force microscope, are described in our previous work (Liu et al. 2019; Nelsen et al. 2019). Beaded cantilevers were generated by first drying 6.0 μm carboxylate beads onto a coverslip (17141-5, Polysciences, Inc); a small amount of UV-curable glue (NOA81, Norland Products) was spread onto the cover slip. A cantilever (Arrow TL1, Nanoworld) was mounted onto the AFM head (Ayslum Research MFP3D, Oxford Instruments), which was lowered over the aforementioned coverslip. Using the manual height adjustment on the AFM, the cantilever was lowered first into the glue and then overtop of a bead. A UV flashlight was used to cure the glue for one minute; the cantilever was removed and set to cure for an additional five minutes. Once beaded, cantilevers were calibrated in media using the thermal tune method (Gavara 2017); the nominal spring constant was 0.03 N/m.

Cells prepared as described above were place onto the AFM-LS system in a custom, 3D-printed mount (uploaded as Thingiverse 2035546). An objective lens heater (Hk-100, Thorlabs, Inc, USA) with a PIV controller (TC200, Thorlabs, Inc.) and a heated scanning stage (900.062 MFP3D Scanner, Oxford Instruments) were used to keep the sample at 37°C. The AFM headed with a calibrated cantilever was placed atop the sample and the cantilever was lowered over a cell of interest. Side-view imaging is achieved by placing a small (180 μm) mirror (8531-607-1, Precision Optics Corporation) adjacent to a cell of interest and raising the objective lens (UplanSAPO 60x/1.2 W, Olympus) until the image plane intersects the mirror. Details regarding mirror alignment and production are given in our previous work (Nelsen et al. 2019). With the AFM in place, the mirror is placed next to the cell of interest such that the cantilever sits between the mirror and the cell. A vertical light sheet propagates out of the objective lens and an electrically tunable lens (ETL) was used to ensure the waist of the light sheet was in the cell. A second ETL is used in the detection path to dynamically adjust focus without moving the objective lens (Liu et al. 2019).

Force curves were taken at a loading rate of 1 μm/s unless otherwise stated. The trigger point for the z-piezo movement was set such that the nucleus was compressed to approximately 2 μm. The z-piezo was then fixed in a closed-loop feedback mode for 60 s, after which the AFM retraced and continue recording data for an additional 15 s. Data from the AFM was recorded at a bandwidth of 2 kHz. A square wave from the AFM was sent to a DAQ board (PCIe-6323, National Instruments) which was used to synchronize both the camera (ORCAFlash4.0 V3, Hamamatsu) and laser light (OBIS-561-150-LS and OBIS-488-150-LS, Edmund Optics). Unless otherwise stated, each channel (488 nm and 561 nm) had an exposure time of 100 ms and 25 ms was taken between each frame resulting in a two-color frame acquisition rate of 4 Hz. Custom software was designed for the synchronization process and image acquisition.

### Nuclear morphology extraction and curvature analysis

All nuclear morphology extraction was performed in FIJI (Schindelin et al. 2012) using side-view fluorescence images of H2B. A rolling-ball background subtraction was performed with a radius dependent on the size of the nucleus (~50 – 150 pixels). A Gaussian blur was then performed with a kernel size of 2 pixels based upon the full-width half max (FWHM) of the system’s point spread function (PSF). The FeatureJ Edges plugin (http://imagescience.org/meijering/software/featurej/) was used to determine the outline of the nucleus during compression; this outline was thresholded to generate a binary image. The binary outline was then dilated several pixels (2 – 5 pixels, depending on the initial image quality) to form a continuous boundary, after which the boundary was filled to generate a mask. The mask was eroded by the same amount as the initial dilation to form the final mask of the nucleus. FIJI’s “analyze particles” feature was used to extract the cross-sectional area and perimeter of the nucleus throughout the AFM compression.

All curvature analysis was performed in Mathematica 11.2 (https://github.com/alihashmiii/curvatureMeasure). Masks of nuclei were imported and the mask boundary was discretized such that each discrete point was separated from the next by approximately 250 nm based upon the FWHM of the PSF of our system. For each point on the perimeter, a circle was fit to that point and the adjacent points within one fourth of the circumference of the AFM bead on either side. Curvature was defined to be the inverse of the radius of the fitted circle; a curvature of 0 represents a flat line. To dynamically track the maximum curvature at the site of indentation, a Gaussian curve was fit to the curvature versus boundary point data in the region where the nucleus was indented

### Treatments of SKOV3 cells

For treatment with Trichostatin-A (TSA), TSA was dissolved to 10 mM in DMSO, and then serially diluted in PBS to 4 μM on day of treatment. 10 μL of a 4 μM solution in PBS was then added to 190 μL of media in 10 mm cloning cylinders, for a final concentration of 200 nM. Experiments were carried out 24-28 hours after drug addition. The 2 × 10^−5^ dilution of DMSO, giving 0.002% v/v final concentration was judged to be insignificant to the TSA effect. A full description of our knockdown of Lamin A/C is available in our previous work (Stephens et al. 2017) Briefly, DNA for the interfering RNA was transfected using Fugene HD. The media was changed regularly after the first 2-day treatment. Cells were plated on days 3, 4, and 5, and used on days 4, 5, and 6 respectively. Two cells were excluded from the lamin A/C sample because they showed sever plastic damage, a phenomenon previously observed in the literature (Pajerowski et al. 2007; Cao et al. 2016) but not present in any other cell of the study (Figure S7).

### Immunofluorescence

To test whether the TSA treatment was effective at inhibiting histone deactylation, we plated cells as described and treated half with TSA as described, and half with a sham pipetting of PBS. At 24 hours, cells were fixed in 4% formaldehyde, permeabilized with .25% Triton x-100, washed and incubated with rabbit monoclonal antibody to acetyl histone H3 (Acetyl-Histone H3 (Lys9) (C5B11) Rabbit mAb #9649, Cell Signaling Technology), 1/400 dilution with 1 mg/mL BSA as blocker, overnight. After primary antibody incubation, cells were washed 3 times, and incubated with Alexafluor 488 goat anti-rabbit IgG (Invitrogen), and 1/1000 dilution of Hoechst 33342 DNA stain for 1 hour. After washing, cells were imaged at 150 ms exposure with 405 nm and 488 nm excitation light (Objective lens: Plan Apo 60x/1.20 W, Nikon) (Figure S4A and B).

To test whether the knockdown was successful, we fixed and stained parallel samples with Lamin A/C Antibody (E-1) (sc-376248, Santa Cruz Biotechnology) as described above, but used a Goat anti Mouse secondary antibody (Alexafluor 568, Invitrogen). Cells plated in parallel were stained with the same solutions and were imaged at 100 ms exposure with 488 nm and 568 nm excitation light (Objective lens: UPlanFL N 40x/1.3 Oil, Olympus) (Figure S5A).

After the collection of immunofluorescence images, nuclei were manually segmented in FIJI (Schindelin et al. 2012). For the TSA verification, mean intensity of the H3K9ac marker was calculated for all nuclei (n = 43 for WT, n = 41 for TSA). A T Test shows a significant relative increase in H3K9ac of approximately 250% for TSA treated cells as compared to WT cells (Figure S4C). For the Lamin A/C KD verification, mean intensity of lamin was quantified for cells expressing the GFP reporter in LA/C KD nuclei (n = 27) and for all WT nuclei (n = 58). A T Test shows a significant relative reduction in lamin expression for LA/C KD nuclei of approximately 40% as compared to WT nuclei (Figure S5B)

### Finite element analysis

All finite element analysis (FEA) modeling was performed in COMSOL Multiphysics 5.2a, the full model is available upon request. The geometry was defined in 2D under axisymmetric assumptions. The AFM tip was modeled to be an elastic sphere of radius 3 μm with an elastic modulus of 3.5 GPa and Poisson ratio of 0.3. The nucleus was modeled to be an elastic ellipsoid with a long axis radius of 8.5 μm and a short axis radius of 3 μm. The bottom 1.5 μm of the ellipsoid was truncated and set to be a fixed constraint. The nucleus had a Poisson ratio of 0.3 and an elastic modulus varied about 1 kPa. The nucleus was wrapped in a thin elastic layer governed by a total spring constant which varied about 10 mN/m. The thin elastic layer also changes the boundary condition between the nucleus and AFM tip such that forces on either side of the boundary are equal in magnitude and opposite in direction, but the displacements on the either side of the boundary are no longer coupled. To simulate indentation, the AFM tip was incrementally stepped a total distance of 2 μm in steps of 0.1 μm. At each step, the MUMPS solver was used to solve for the displacement and stress along the mesh. A surface integral on the AFM tip provided the reactionary force at each indentation step; nuclear morphology was also extracted at each indentation step. All analysis was performed under that assumption of quasistatic behavior; that is, no time-dependence was accounted for in our FEA model. The equation governing the behavior of elastic solids is given by

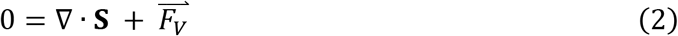

where **S** is the stress tensor and 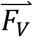 is the volume force. The equation governing the thin elastic layer is given by

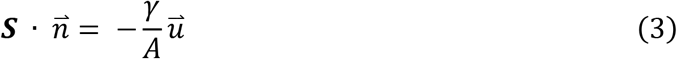

where ***S*** is the stress tensor, 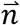 is the vector normal to the surface, *γ* is the stretch modulus, *A* is the contact area, and 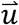 is the displacement field.

## Abbreviations

NCSA: Nuclear Cross-Sectional Area
NP: Nuclear Perimeter
AFM: Atomic Force Microscopy
AFM-LS: Combined Atomic Force Microscopy and Light Sheet Imaging System
WT: Wild Type
TSA: Trichostatin A
LA/C KD: Lamin A/C Knock Down
FEA: Finite Element Analysis

## Acknowledgments

We thank the Goldman Lab (Northwestern University Feinberg School of Medicine) for providing us with the siRNA plasmid to knockdown lamin A/C levels. C.M.H. is supported by the NSF GRFP (DGE-1650116) and the Caroline H. and Thomas Royster Fellowship. M.K is supported by NIH 5T32GM008570-19. A.D.S is supported by Pathway to Independence Award NIHGMS K99GM123195. E.T.O, M.R.F, and S.R are supported by NIH and NSF (NSF/NIGMS 1361375) as well as NIH (NIBIB P41-EB002025).

