## Supplementary Figures for "Correlating nuclear morphology and external force with combined atomic force microscopy and light sheet imaging separates roles of chromatin and lamin A/C in nuclear mechanics"

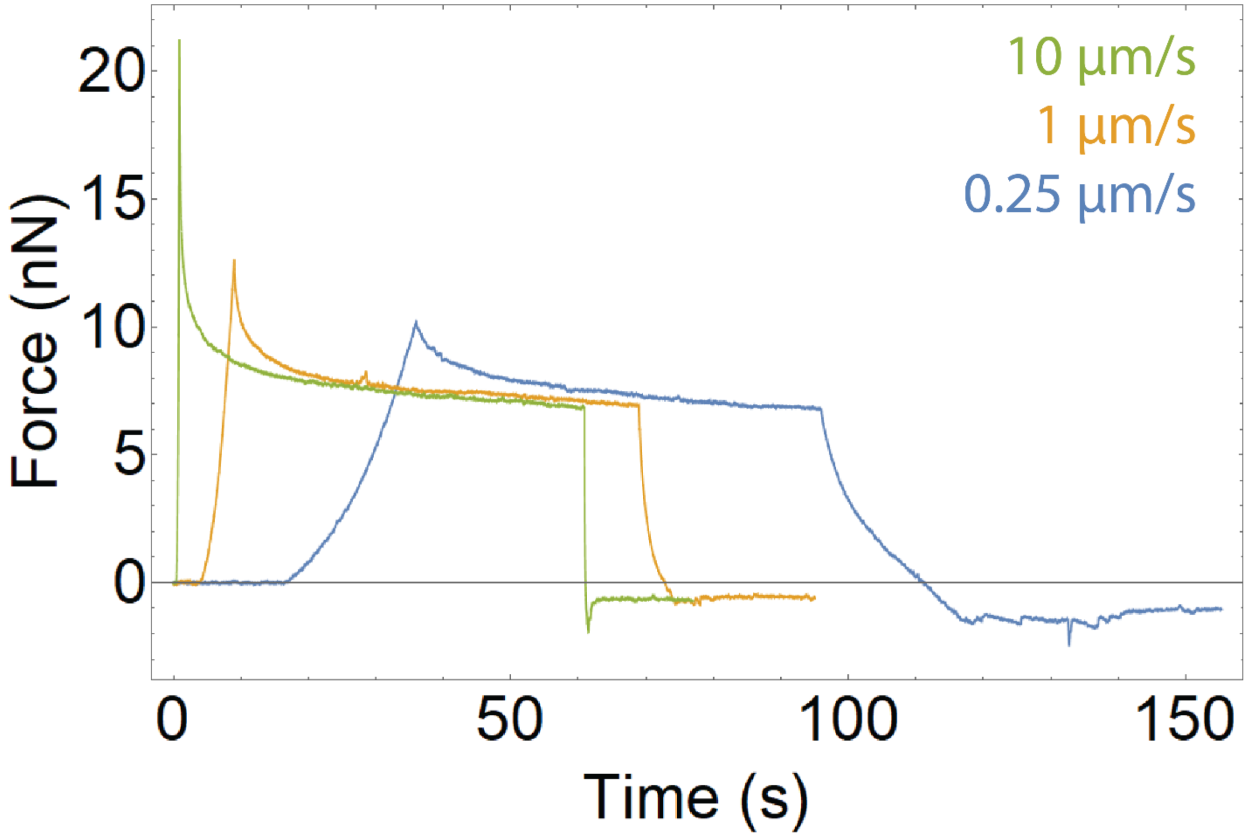


Figure S1: Force versus time plot for three different AFM indentation and retraction rates (10 μm/s – Green, 1 μm/s – Orange, 0.25 μm/s – Blue). All AFM parameters asides from the indentation and retraction rate remained constant. For each compression, the cantilever was lowered a prescribed distance at the given rate, and then the z piezo was held fixed for 60 seconds. The cantilever then retracted at the same rate and the z piezo was help fixed for 15 seconds. Viscous relaxation is present for all rates of indenation, as seen by the decay during the 60 second dwell.

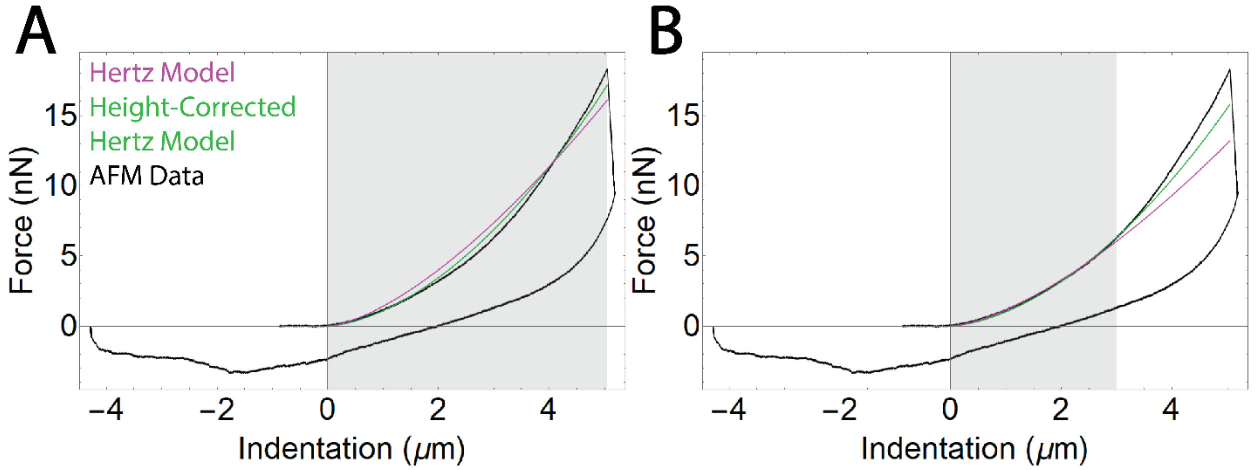


Figure S2: Force versus indentation plots showing the fits of both the Hertz model (Purple) and the height-corrected Hertz model (Green) over the entire indentation (A) and over the first 3 µm of indentation (B). The gray region represents the portion of data to which the model was fit. Fitting to the entire indentation shows that neither model accurately represents the data set. Fitting to the early portion of the indentation shows that both models are applicable for low indentations, but underestimate the force response at high indentations. This is suggestive that an additional mechanism for force response is needed to match the experimental data.

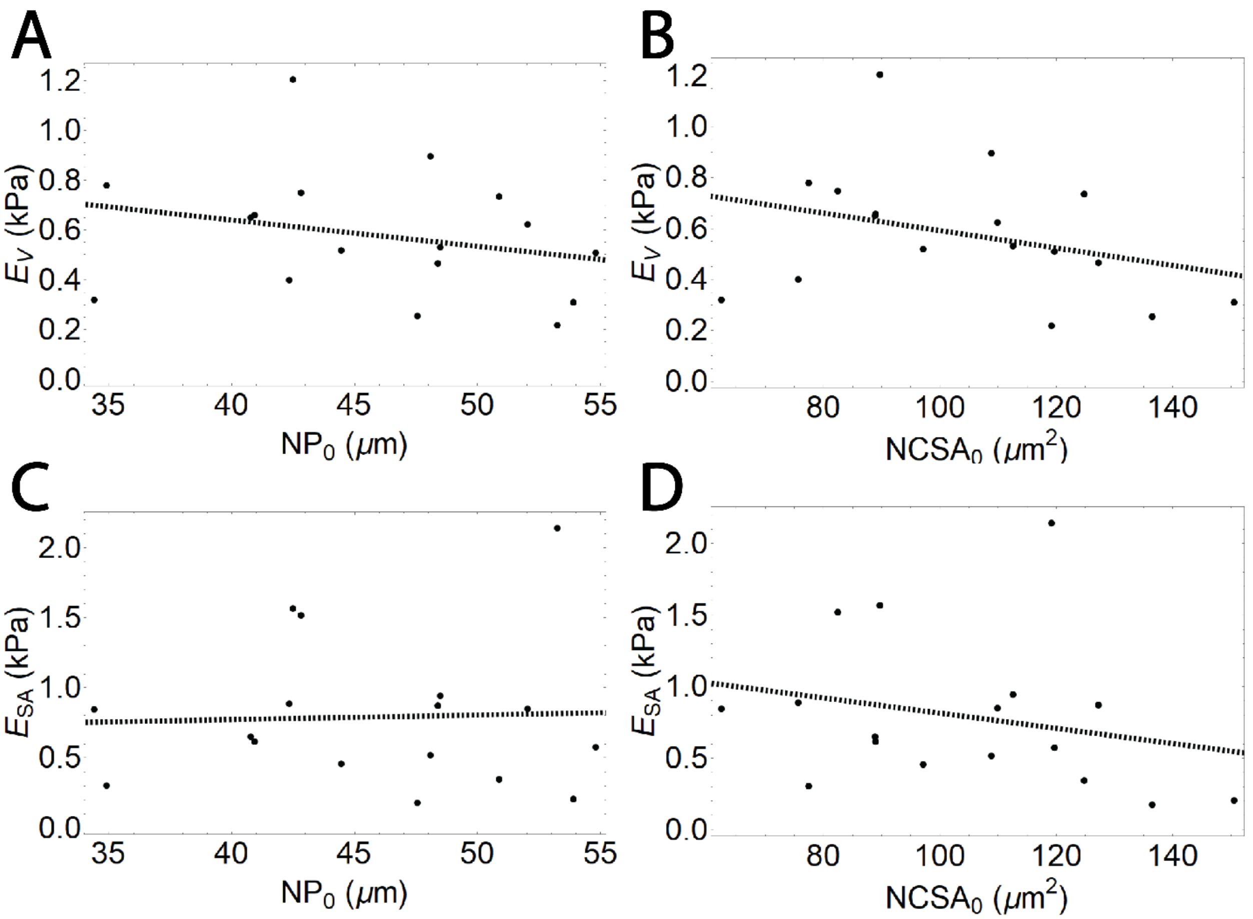


Figure S3: There is no significant correlation between initial nuclear morphology and $E_V$ or $E_{SA}$. Plots of (A) $E_V$ versus $NP_0$, (B) $E_V$ versus $NCSA_0$, (C) $E_{SA}$ versus $NP_0$, (D)$E_{SA}$ versus $NCSA_0$, as well as best fit lines (dashed). A Pearson's correlation test for each relationship shows no significant correlation.

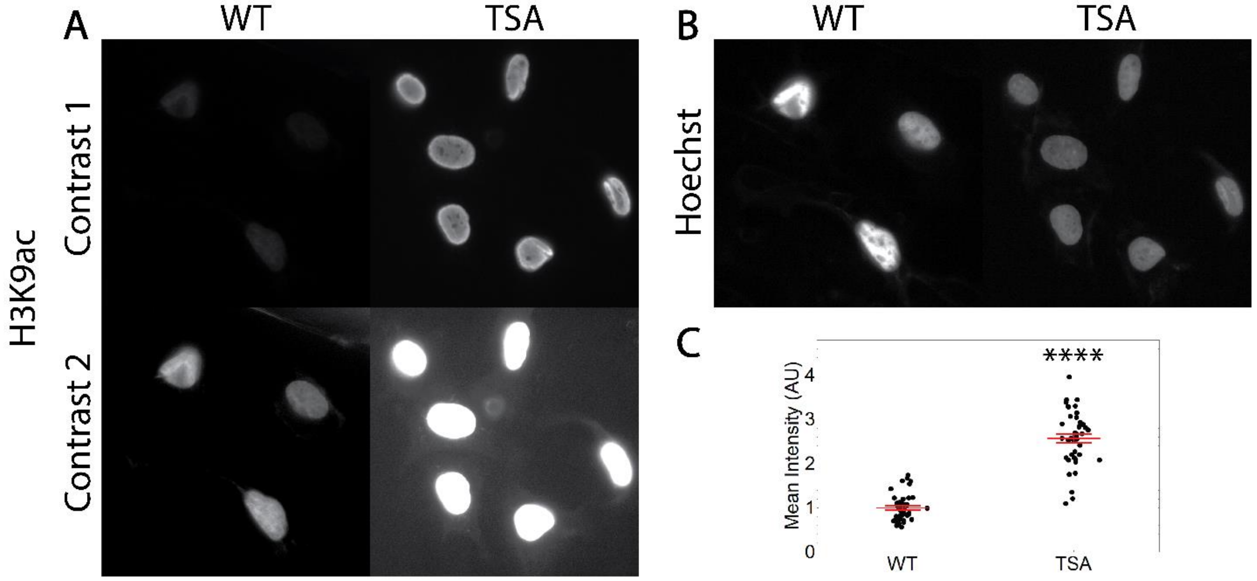


Figure S4: Immunofluorescence validation of TSA treatment. (A) Representative fluorescence images of H3K9ac in both WT and TSA treated cells. Images are shown at two different contrast levels. (B) Representative fluorescence images of Hoechst in both WT and TSA treated cells. (C) Mean intensity of H3K9ac for WT (n = 43) and TSA treated (n = 41) cells shows a significant increase in decondensed chromatin for TSA treated cells. **** represents p < 0.0001 for a T Test. Red lines represent mean and standard error.

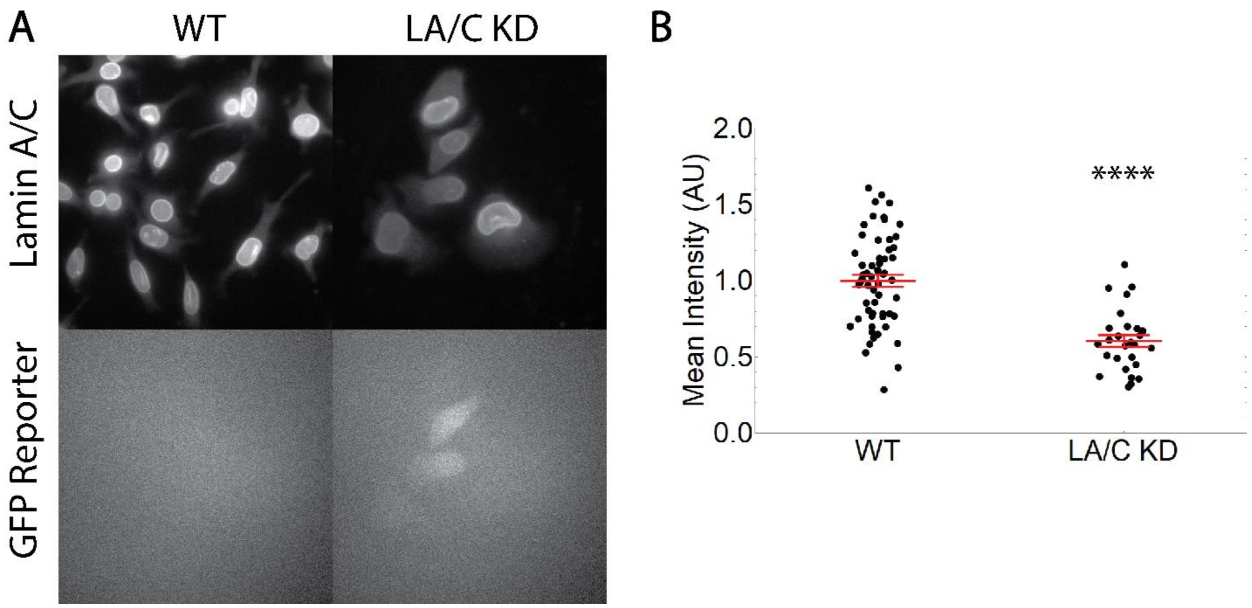


Figure S5: Immunofluorescence validation of lamin A/C knock down. (A) Representative fluorescence images of lamin A/C and the GFP reporter in both WT and LA/C KD cells. (B) Mean intensity of lamin A/C for WT (n = 58) and LA/C KD (n = 27) cells shows a significant decrease in lamin expression for LA/C KD cells. **** represents $p < 0.0001$ for a T Test. Red lines represent mean and standard error.

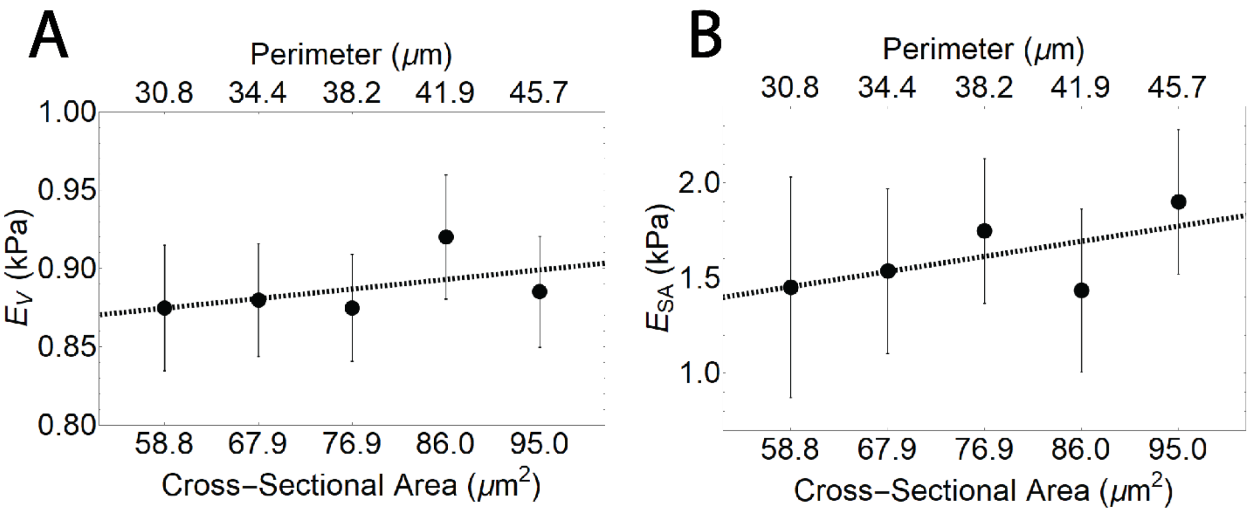


Figure S6: $E_V$ and $E_{SA}$ extracted from FEA model are independent of initial nuclear morphology. Plots of $E_V$ (A) and $E_{SA}$ (B) versus nuclearly morphology as well as best fit lines. A Pearson's correlation test gives no significant correlation between $E_V$ or $E_{SA}$ and initial nuclear morphology.

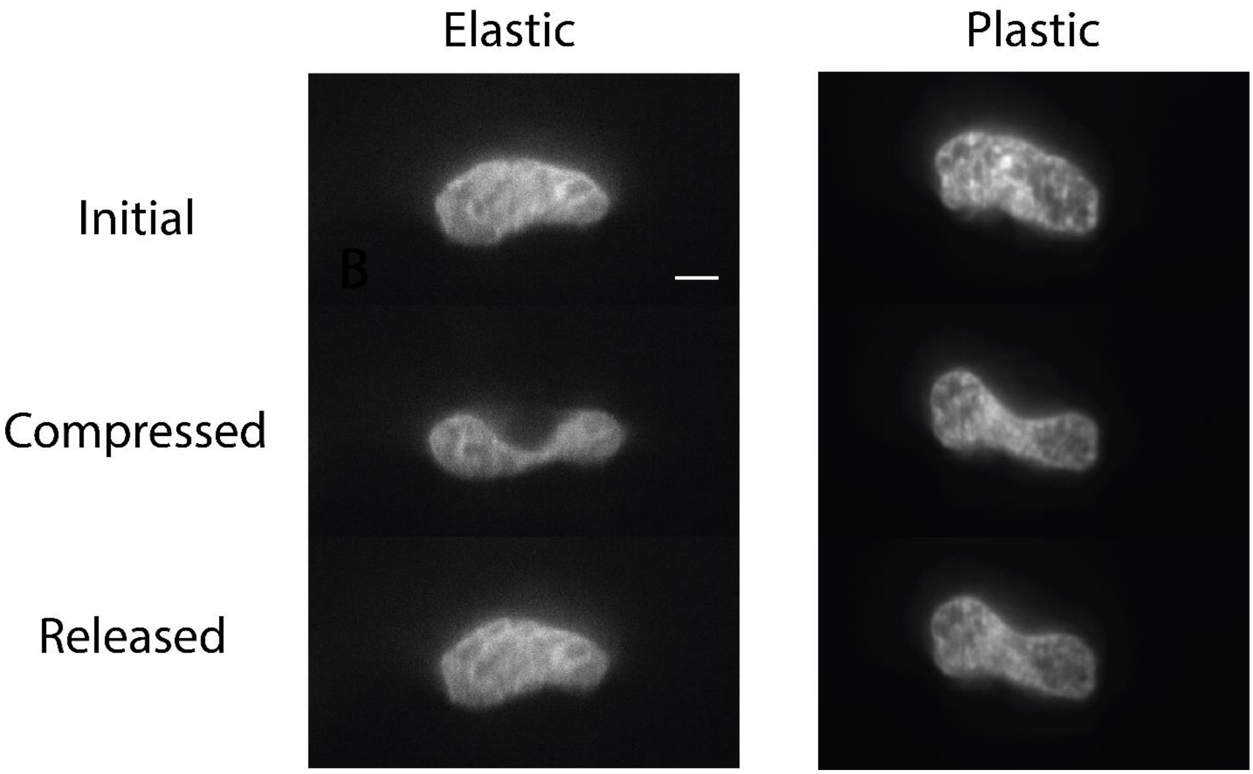


Figure S7: Side-view image series for AFM compression of live SKOV3 stably expressing halotagged H2B labeled with Janlia Fluor 549. These cells were compressed 6 days post-transfection of siRNA to halt production lamin A/C. One nucleus (left) shows elastic behavior, while the second nucleus (right) shows plastic damage. Scale bar = 5 µm.

**Supplementary Movie Captions**

Movie S1: Combined side-view light sheet microscopy and atomic force microscopy. (Left) Force versus time series during AFM indentation. Red line indicates current frame. (Right) Side-view images of a live SKOV3 cell under compression. Green – halotagged-H2B labeled with Janelia Fluor 549. Magenta – snaptagged-Kras Tail labeled with Janelia Fluor 503. Scale bar = 5 µm.

Movie S2: Dynamic curvature analysis curing AFM compression. (Left) Nuclear curvature versus position along perimeter of the nucleus during AFM compression. Note the formation of a peak at approximately 10 µm throughout the compression. (Right) Side-view images of a live SKOV3 cell under compression. Green – halotagged-H2B labeled with Janelia Fluor 549. Magenta – snaptagged-Kras Tail labeled with Janelia Fluor 503. Scale bar = 5 µm.

Movie S3: Finite element analysis simulation of AFM compression. Grid lines indicate distance in µm. Color map indicates von Mises stress in Pa.
